# Fecal Dysbiosis and Inflammation in Intestinal-Specific Cftr Knockout Mice on Regimens Preventing Intestinal Obstruction

**DOI:** 10.1101/2023.07.24.550378

**Authors:** Sarah M. Young, Rowena A. Woode, Estela Williams, Aaron Ericsson, Lane L. Clarke

## Abstract

Chronic intestinal inflammation is a poorly understood manifestation of Cystic Fibrosis (CF), which may be refractory to ion channel CFTR modulator therapy. People with CF exhibit intestinal dysbiosis which has potential for stimulating intestinal and systemic inflammation. CFTR is expressed in organ epithelia and in the leukocyte population. Here, we investigate the contribution of intestinal epithelial-specific loss of Cftr (iCftr KO) to dysbiosis and inflammation in mice treated with either of two anti-obstructive dietary regimens necessary to maintain CF mouse models (PEG laxative or a liquid diet, LiqD). Feces collected from iCftr KO mice and their wildtype (WT) sex-matched littermates were used to measure fecal calprotectin and to perform 16S rRNA sequencing to characterize the gut microbiome. Fecal calprotectin was elevated in iCftr KO relative to WT samples of mice consuming either PEG or LiqD. PEG iCftr KO mice did not show a change in α-diversity versus WT but demonstrated a significant difference in microbial composition (β-diversity) with increases in phylum *Proteobacteria*, family *Peptostreptococcaceae*, four genera of *Clostridia* including *C. innocuum*, and mucolytic genus *Akkermansia*. Fecal microbiome analysis of LiqD iCftr KO mice showed both decreased α-diversity and differences in microbial composition with increases in *Proteobacteria* family *Enterobacteriaceae*, *Firmicutes* families *Clostridiaceae* and *Peptostreptococcaceae*, and enrichment of *Clostridium perfringens*, *C. innocuum*, *C. difficile*, mucolytic *Ruminococcus gnavus*, and reduction of *Akkermansia*. It was concluded that epithelial-specific loss of Cftr is a major driver of CF intestinal dysbiosis and inflammation with significant similarities to previous studies of global Cftr KO mice.

**New and noteworthy:** Chronic intestinal inflammation is a manifestation of cystic fibrosis (CF), a disease caused by loss of the anion channel CFTR that is expressed in many tissues. This study shows that intestinal epithelial cell-specific loss of CFTR (iCftr KO) in mice is sufficient to induce intestinal dysbiosis and inflammation. Studies were performed on mice consuming either dietary regimen (PEG laxative or liquid diet) routinely used to prevent obstruction in CF mice.

## Introduction

Cystic fibrosis (CF) is a mono-genic inheritable disease that occurs in approximately 31,000 individuals in the United States and 162,000 individuals globally(1, 2). Loss of function mutations in the cystic fibrosis transmembrane conductance regulator (CFTR) gene leads to multi-organ disease centered around polymeric mucin-producing epithelial surfaces(3). In the CF intestine, the occurrence of chronic low-grade inflammation is a poorly understood manifestation characterized by pro-inflammatory cytokine expression, increased mucosal IgA and IgM immunoglobulins, and increased neutrophil chemotaxis(4). Imaging modalities including capsule endoscopy in CF children and adults have demonstrated gross pathological changes to the small intestinal mucosa suggestive of inflammation including erythema, mucosal ulceration and breaking without clinical disease(5). Microbial dysbiosis, another reported manifestation in children and adults with CF(6–8), has been shown to be associated with intestinal inflammation in patients affected by inflammatory bowel-like diseases(9). Recognized patterns in intestinal microbial changes in children with CF include a reduction in taxa diversity and richness, and an enrichment of members of the *Proteobacteria* phylum including *Escherichia coli*(7). The gut microbiota disturbances commonly seen correlate with recognized microbial signatures for Crohn’s Disease (e.g., enrichment of *Escherichia* and *Ruminococcus* with overt decrease in taxa richness and diversity)(9). Various alterations to the intestinal intraluminal environment including mucus accumulation(10), decreased pH(11), potential for reduced Paneth cell anti-microbial granule dissolution(12) and a decrease in gastrointestinal transit time(13) may impact bacterial colonization and community dynamics in CF disease.

The *Cftr*^tm1UNC^ (null mutation, pan-Cftr KO) mouse model of CF has been shown to recapitulate intestinal inflammation and dysbiosis as observed in human patients(14–17). Chronic low-grade inflammation throughout life, as reported in people with CF (pwCF), has been described in pan-Cftr KO mice and has been associated with increased risk of gastrointestinal neoplasia in both cases(18–22). The advantage of the pan-Cftr KO mouse as a research model for inflammation and dysbiosis in pwCF is the lack of overt lung and pancreatic disease(23). Further, antibiotic or enzyme replacement therapies are not required, which greatly improves the clarity of interpretation for intestinal and microbiota differences in CF mice as compared to wild-type (WT) mice. However, one disadvantage of intestinal disease in pan-Cftr KO mice is obstipation and consequent mortality due to tenacious mucofeculent impactions in the ileum leading to peritonitis(24). To prevent the development of this condition, post-weaning mice are typically maintained on one of two dietary regimens. In the first case, WT and pan-Cftr KO mice are fed standard lab chow but polyethylene glycol 3350 m. wt. (PEG) laxative and electrolytes are added to the drinking water (PEG regimen). The other dietary regimen is the provision of a nutritionally-complete, liquid diet (LiqD regimen) (Peptamen®, Nestles). Both regimens have been shown to enable researchers to produce pan-Cftr KO mice with adult murine life-spans(25, 26).

There is increasing biomedical research interest in CF intestinal disease and consequent changes of the intestinal microbiota. With the advent of highly effective modulators of mutant CFTR, e.g., Trikafta®, life expectancy of pwCF is expected to significantly increase along with an increasing prevalence of CF intestinal disease(1). Trikafta® in an observational clinical trial (PROMISE GI) of pwCF indicated only a small benefit with regard to most gastrointestinal symptoms after 6 months(27). Based on earlier reports of dysbiosis in CF mice(12, 14-17, 28), studies have begun dissecting aspects of the CF intestinal microbiome. Germ-free pan-Cftr KO mice were found to develop both systemic inflammation and dysbiosis when gavaged with fecal microbial transplants from specific-pathogen-free inbred WT mice, indicating that CF intestinal pathogenesis is a direct consequence of Cftr loss(14). Studies of the mucosa-attached microbiota of the Cftr KO small intestine found increased abundance of pro-inflammatory *Escherichia* and depletion of beneficial secondary bile acid producing bacteria(17). It is also known that CFTR is expressed in both epithelial and non-epithelial tissues such as smooth muscle and leukocytes(29, 30). In a recent investigation of the CF leukocyte contribution to intestinal dysbiosis, Wang and colleagues used Cftr exon 11 floxed mice(31) to generate mice that lacked Cftr expression in the myeloid, macrophage or neutrophil populations for comparison to WT and pan-Cftr KO mice(32). The pan-Cftr KO mice showed the most dramatic decreases in alpha diversity (microbial richness, evenness and phylogenetic diversity). Changes in the evenness (Shannon Index) and phylogenetic diversity (Faith’s PD) were also apparent in the myeloid Cftr KO, validating an innate immune deficiency in CF myeloid cells. With regard to the microbial similarities as measured by principal component analysis (Bray-Curtis dissimilarity), the pan-Cftr KO mice were most dissimilar among the five genotypes in both small and large intestinal/fecal samples.

In the present study, we evaluate fecal calprotectin as a measure of inflammation and the composition of the fecal microbiota between sex-matched, littermate WT and intestinal-specific *Cftr* knockout B6.Cg-Tg(Vil1-cre)-*Cftr^f10/f10^* mice (iCftr KO) to evaluate intestinal epithelial cell-autonomous consequences of Cftr loss. To the best of our knowledge, the impact of Cftr loss in intestinal epithelial cells alone on microbial colonization in the large intestine has not been reported. As epithelial loss of CFTR causes significant environmental changes in the gut environment, we hypothesized that *Cftr* loss in the intestinal epithelium alone would drive a dysbiotic phenotype. To extend the utility of CF mice for intestinal investigation, fecal calprotectin and characteristics of the microbiome were compared on fecal samples of iCftr KO and WT mice consuming either the PEG or LiqD regimens.

### Materials and methods Ethics statement

This study was performed in accordance with the 8^th^ Edition of the Guide for the Care and Use of Laboratory Animals and approved by the University of Missouri Institutional Animal Care and Use Committee (MU ACUC protocol #9771). All animals were housed and cared for in an Association for Assessment and Accreditation of Laboratory Animal Care (AAALAC) accredited vivarium.

### Mice

B6.Cg-Tg(Vil1-cre)-*Cftr^f10/f10^*, intestinal-specific *Cftr* knockout (i*Cftr* KO) mice were produced as previously described through the crossing of *Cftr^fl10^* conditional *Cftr* mice with mice caring a Villin-Cre transgene which expresses Cre recombinase only in villus and crypt intestinal epithelial cells(33). The mice were backcrossed to the C57BL/6J background for greater than 10 generations. Breeder pairs, offspring, and weaned mice were maintained on standard pelleted laboratory chow (Formulab 5008 Rodent Chow) *ad libitum* and a standard osmotic laxative drinking water preparation containing 10 mM polyethylene glycol 3350, 69 mEq of Na^+^, 5 mEq of K^+^, 11 mEq of HCO_3_ ^-^, 22 mEq of SO ^2-^, and 19 mEq of Cl^-^ per liter reconstituted with deionized water. Upon weaning, all mice were isolated in individual cages for the duration of the study for two reasons. First, iCftr KO mice are smaller than WT mice and do not compete well for access to food in grouped housing. Second, a goal of the study was to assess individual mouse responses on each anti-obstructive dietary regimen to inform researchers using one or the other regimen for studies of CF mice. The experiment design is internally controlled by comparisons between WT and iCftr KO sex-matched littermates. For fecal microbiota studies, adult mice (2-4 months) of both sexes were maintained in clean open-top corncob bedded cages for 14 days and provided either the PEG regimen or the LiqD regimen for two weeks before fecal sample collection. The LiqD regimen was by provision of Peptamen® (Nestle Health Science), a nutritionally complete liquid diet, which was changed daily in cage mounted J-feeders, and tap water. For fecal calprotectin experimentation, WT and iCftr sex-matched littermate pairs were first maintained on PEG regimen for collection of feces and then transitioned to LiqD regiment for 14 days for a second feces collection. All mice were monitored daily for body weight, feed consumption, behavior, and activity.

### Sample Collection

Freshly evacuated fecal pellets were collected from each sex-matched iCftr KO and WT littermate pairs by transferring each mouse individually to a clean plastic container swabbed with 70% ethanol. Each mouse was allowed to defecate normally. Fecal pellets were collected into sterilized 1.5 mL Eppendorf tubes using 70% ethanol-disinfected forceps. Samples were immediately stored at −80^°^ C until DNA extraction or calprotectin measurement was performed. All fecal samples were collected between 7 and 9 a.m., CDT.

### Fecal Calprotectin Measurement

Calprotectin concentration of fecal pellets was measured by ELISA, according to the manufacturer’s protocol (Immundiagnostik AG, Bensheim, Germany). Briefly, fecal pellets and assay reagents were brought to room temperature. Fecal samples were mixed with extraction buffer and centrifuged for 10 minutes at 3000 x g at room temperature (RT). Wells were washed 5 times with wash buffer and 100 µl of sample supernatant and assay standards were added to the wells of the assay plate, covered, and incubated for 1 hour at 37°C. The plate was washed 5 times with wash buffer, and 100 µl of monoclonal anti-calprotectin antibody was added and incubated for 1 hour at 37°C. A second washing procedure was performed and 100 µl of diluted conjugate solution was added to each well and the plate was incubated for another hour at 37°C. After 5 washes, 100 µl of substrate was added to each well, incubated for 15 minutes at RT in the dark. Following the addition of 100 µl of stop solution, absorption was immediately determined using an ELISA plate reader at λ450nm. The fecal calprotectin concentration was calculated from the assay standards and expressed as µg/g feces.

### DNA extraction

A single fecal pellet from each animal was transferred to a sterile 2 mL round-bottom tube containing a stainless-steel bead using sterile wooden toothpicks. DNA extraction was performed using the QIAmp PowerFecal Pro DNA Kit (Qiagen, Venlo, Netherlands), following the manufacturer’s instructions. Mechanical disruption was performed using a TissueLyser II (Qiagen, Venlo Netherlands). DNA was eluted in 100 µL EDTA-free Solution C6. DNA yield was determined via fluorometry (Qubit, Life Technologies, Carlsbad, CA) using the quanti-iT BR dsDNA reagent kit (Invitrogen, Carlsbad, CA). DNA samples were frozen at −80^°^C prior to submission for 16S rRNA sequencing through the University of Missouri Metagenomics Center (MUMC).

### 16S rRNA library preparation and sequencing

Extracted DNA was processed for 16S rRNA sequencing at the University of Missouri Genomics Technology Core Facility. Bacterial 16S rRNA amplicons were constructed using the highly conserved V4 region of the 16S rRNA gene using universal primers (U515F/806R) flanked by Illumina® standard adapter sequences. Oligonucleotide sequences used are available for review at proBase(34). Dual-indexed forward and reverse primers were used, allowing multiple samples to be sequenced per run. Fifty µL reactions containing 100ng of metagenomic DNA, primers (0.2 µM concentration), dNTPs (200 µM), and Phusion high-fidelity DNA polymerase (1U, Thermo Fisher) were used for PCR performance. PCR cycle parameters were 98°C (3 min) + [98°C (15 sec) + 50°C (30 sec) + 72°C (30 sec)] × 25 cycles + 72°C (7 min). Amplicon pools (5 µL/reaction) were combined, thoroughly mixed, and purified by addition of Axygen Axyprep MagPCR clean-up beads to an equal volume of 50 µL of amplicons and incubated for 15 minutes at RT. Products were washed repeatedly with 80% ethanol and the resultant pellet was resuspended in 32.5 µL EB buffer (Qiagen), incubated at RT for two minutes, and placed on the magnetic stand for five minutes. The final amplicon pool was evaluated using the Advanced Analytical Fragment Analyzer automated electrophoresis system, quantified using quant-iT HS dsDNA reagent kits, and diluted according to Illumina’s standard protocol for sequencing on the MiSeq instrument.

### Bioinformatics

Assembly, binning, and annotation of DNA sequences was performed at the University of Missouri Bioinformatics and Analytics Research Core Facility. Primers were designed to match the 5’ ends of the forward and reverse reads. Cutadapt (version 2.6; https://github.com/marcelm/cutadapt) was used to remove the primer from the 5’ end of the forward read(35). If found, the reverse complement of the primer to the reverse read was then removed from the forward read as were all bases downstream. Thus, a forward read could be trimmed at both ends if the insert was shorter than the amplicon length. The same approach was used on the reverse read, but with the primers in the opposite roles. Read pairs were rejected if one read or the other did not match a 5’ primer, and an error-rate of 0.1 was allowed. Two passes were made over each read to ensure removal of the second primer. A minimal overlap of three bp with the 3’ end of the primer sequence was required for removal.

The QIIME2 DADA2 plugin (version 1.10.0)® was used to denoise, de-replicate, and count amplicon sequence variants (ASVs) incorporating the following parameters: 1) forward and reverse reads were truncated to 150 bases, 2) forward and reverse reads with number of expected errors higher than 2.0 were discarded, and 3) chimeras were detected using the “consensus” method and removed. R version 3.5.1 and Biom version 2.1.7 were used in QIIME2. Taxonomies were assigned to final sequences using the Silva.v132 database and the classify-sklearn procedure.

Principal coordinate analysis (PCoA) was performed on ¼ root-transformed ASV relative abundance data for each sample, and alpha-diversity indices were calculated using the Past 4.03 software(36) package. A heatmap of log-transformed relative abundance data was arranged via hierarchical clustering using Metaboanalyst 5.0(37).

### Statistical analysis

Differences in fecal microbial community composition were tested via one-way permutational multivariate analysis of variance (PERMANOVA) by weighted Bray Curtis dissimilarity distances using Past 4.03. Differences in Shannon_H diversity and Chao-1 abundance indices were determined through unpaired t-test. Relative abundance of bacteria of interest were first calculated by average sequence reads per 50915 total reads/sample, and unpaired two-tailed T-tests were performed comparing WT to iCftr KO. All unpaired t-test statistical analyses were made using SigmaPlot® 14.0 software. Multiple-testing correction was performed using the Benjamini-Hochberg procedure.

## Results

### The iCftr KO mouse exhibits increased fecal calprotectin with both PEG and LiqD dietary regimens

It was previously shown that small intestinal expression of genes involved in innate defenses, mast cell and neutrophil infiltration are elevated in pan-Cftr KO mice on LiqD, but are reduced in most cases to WT levels in Cftr KO mice receiving the PEG regimen(38). To evaluate intestinal inflammation in iCftr KO mice on the PEG and LiqD regimens, the feces were collected from mice on either regimen for the measurement of fecal calprotectin. Fecal calprotectin is a known biomarker of intestinal inflammation in human IBD cases and recently has been shown to accurately detect low levels of intestinal inflammation(39). As shown in Fig. 1A and 1B, fecal calprotectin was significantly increased in iCftr KO mice as compared to WT on both the PEG regimen and the LiqD. The iCftr KO mice showed greater variability in both cases, especially when consuming the LiqD diet. A comparison of fecal calprotectin of iCftr KO mice on LiqD versus PEG regimen trended toward an increase (*p* = 0.0525). Interestingly, WT fecal calprotectin was significantly different between mice on the LiqD regimen as compared to those on the PEG regimen (*p* = 0.009).

**Figure 1.**
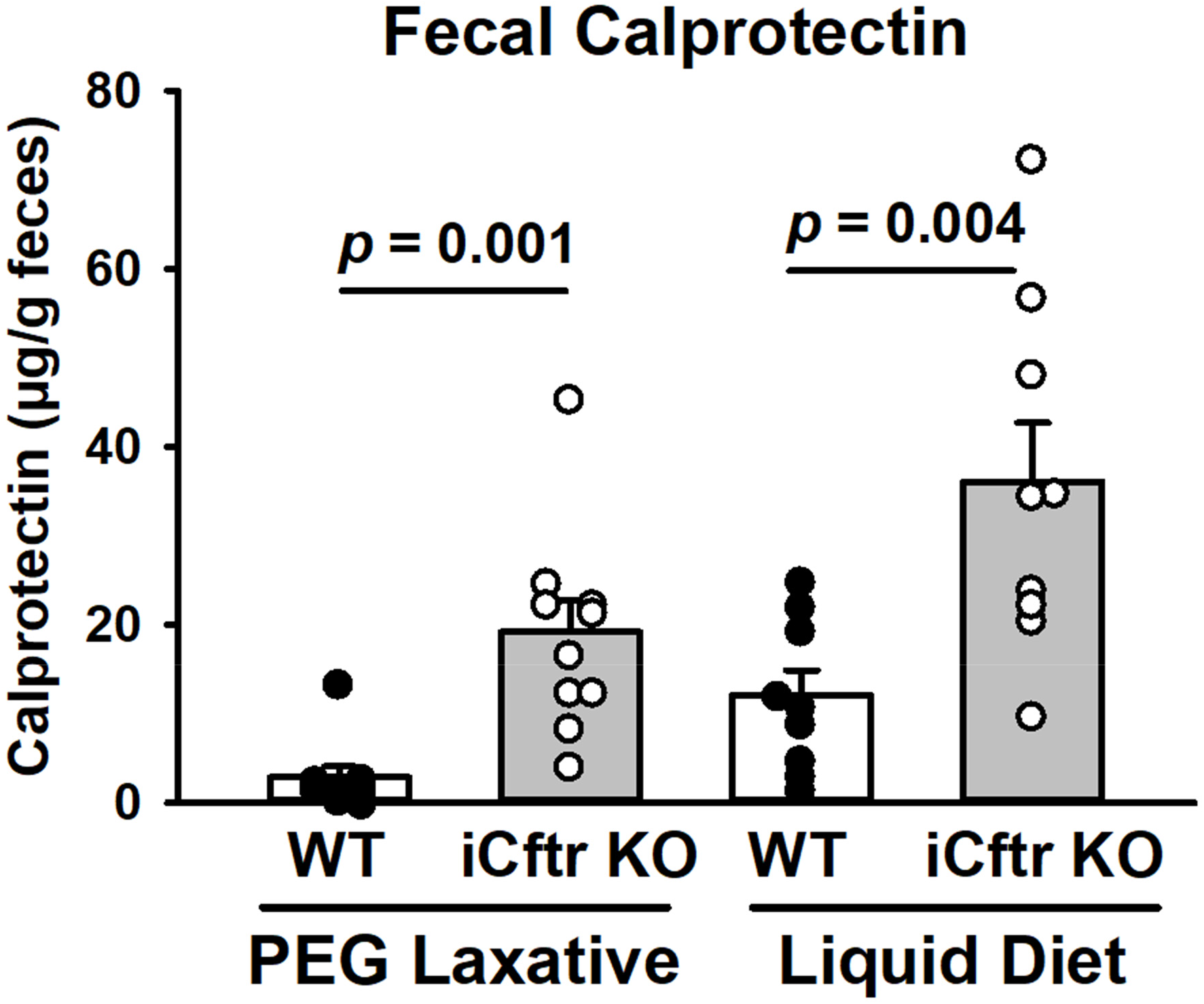
Comparison of fecal calprotectin in WT and iCftr KO mice receiving PEG laxative (left) or liquid diet (right). n = 9 WT and Cftr KO sex-matched littermate pairs, 5 Male, 4 Female. Data represented as individual mouse values (WT – filled circles, iCftr KO – open circles) and mean ± SEM (WT – white bars, iCftr KO – gray bars). Data analyzed using Student’s t-test, p values are shown.

### iCftr KO mice on PEG regimen display altered beta-diversity of bacterial species without significant changes in Shannon diversity and Chao-1 abundance indices

Previous studies have shown that the isolated effect of Cftr ablation in the myeloid immune cell population affects the intestinal microbiome(32). For comparison, we evaluated the isolated effect of intestinal-specific loss of Cftr on the intestinal microbiome. Previous studies of the two anti-obstructive regimens have shown a lower intestinal bacterial load in the PEG versus the LiqD regimen(38). Paralleling this observation, the present study found that the total OTU read counts were significantly less in the feces of PEG-consuming mice versus LiqD-consuming mice (PEG WT = 52,044 ± 1,402 vs. LiqD WT = 78,090 ± 3,188, *p* = 0.000001 and PEG iCftr KO = 54,900 ± 1,866 vs. LiqD iCftr KO = 78,564 ± 5,007, *p* = 0.000244). No significant differences were found for read counts between the iCftr KO and WT feces on either dietary regimen.

A 16S rRNA sequencing analysis of fecal samples from iCftr KO mice and WT littermates maintained on the PEG regimen was used to assess the microbiome. A significant effect of sex on the microbiome was not found, so data from six male and four female iCftr KO mice were combined for cumulative data, as were their corresponding sex-matched WT littermates. As shown in Figure 2A, the relative abundance of bacterial phyla in iCftr KO, as compared to WT, revealed significant increases in *Actinobacteriota* (*Actinomycetota*) (*p* < 0.01), *Verrucomicrobiota* (*p* < 0.05) and *Proteobacteria* (*Pseudomonadota*) (*p* < 0.05), and a significant reduction of phyla *Bacteroidota* and *Desulfobacterota* (*p* < 0.05). At the family level (Figure 2B), *Bifidobacteriaceae*, within the Actinobacteriota phylum, was strongly increased in the iCftr KO feces as compared to that of WT. *Verrucomicrobiota* phylum is dominated by the *Akkermansiaceae* family, which was significantly increased in the iCftr KO feces. In the *Proteobacteria* phylum, the iCftr KO samples showed an increase in the family *Sutterellaceae* of the order *Burkholderiales*. A decrease in the *Bacteroidota* phylum in the iCftr KO feces was matched with a strongly significant decrease in family *Muribaculaceae* but included an increase in the family *Bacteroidaceae* relative to WT specimens. The phylum *Firmicutes* was unchanged between genotypes. However, there were decreases in the families *Erysipelotrichaceae* and *Ocillospiraceae*, and increases in the family *Peptostreptococcaceae*.

**Figure 2.**
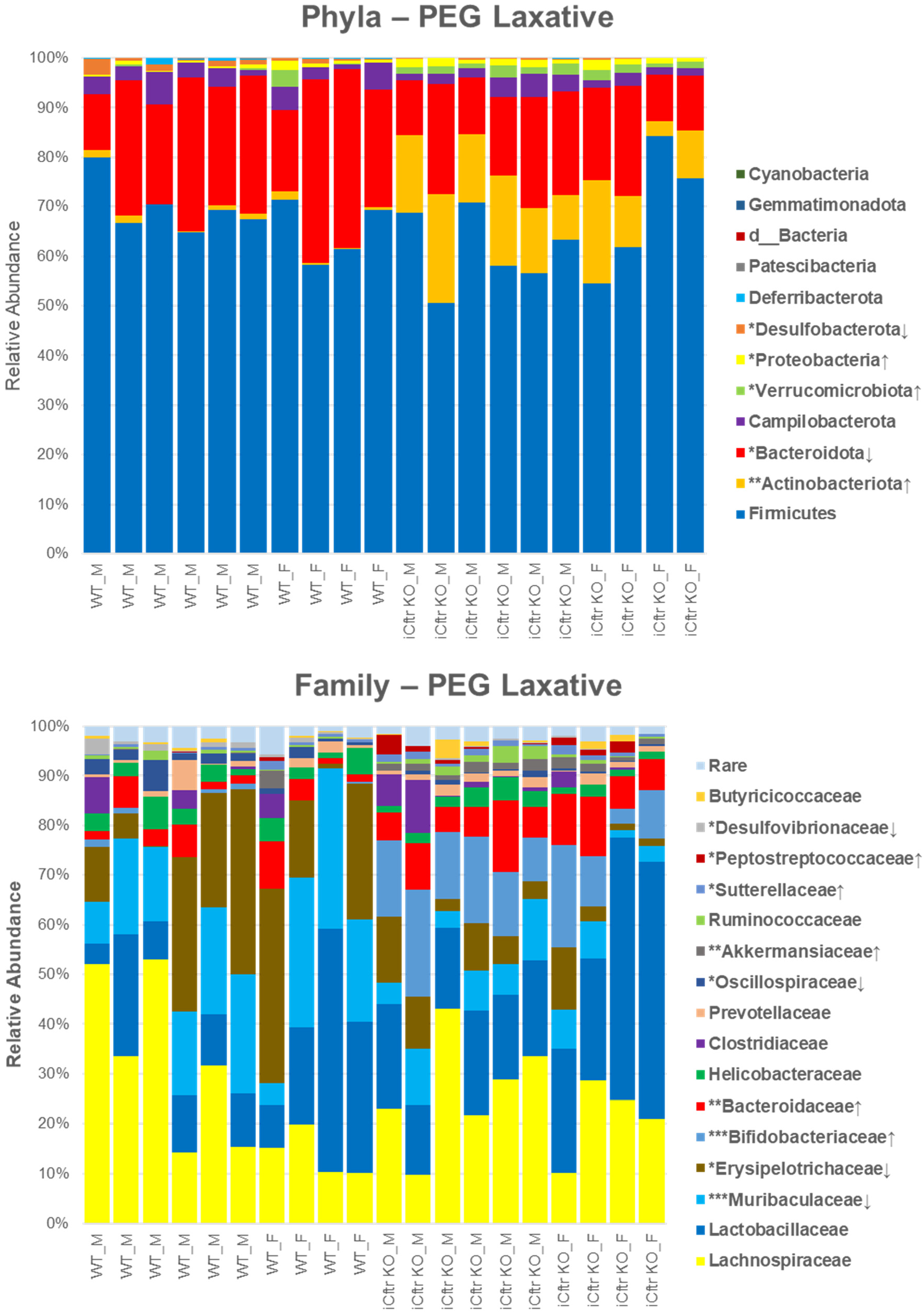
Bacterial taxa of fecal microbiome from WT and iCftr KO sex-matched littermate mice receiving PEG laxative regimen. Top. Stacked bar chart showing taxonomic relative abundance distribution of fecal microbial communities at the phylum level of individual WT and iCftr KO mice receiving PEG laxative. Bottom. Stacked bar chart showing taxonomic relative abundance distribution of fecal microbial communities at the family level of WT and iCftr KO and WT receiving PEG laxative. Samples are grouped by genotype and sex at the x-axis. N = 10 WT and iCftr KO sex-matched littermate paired mice, 6 male, 4 female. Colored squares to the right of the charts indicate phylum or family identity. D = domain. ↑ = significantly increased relative to WT, ↓ = significantly decreased relative to WT. ***p-value<0.001, **p-value<0.01, *p-value<0.05 via unpaired Student’s t-test.

Differences in α-diversity in the fecal microbiome between iCftr KO and WT mice were evaluated using the Shannon_H diversity index (Figure 3A) and Chao-1 index (Figure 3B). No significant difference between the two genotypes was found with either index, indicating similar richness and evenness of the microbiota. In contrast, comparison of β-diversity by PCoA using Bray-Curtis dissimilarity distances revealed significant clustering of samples from i*Cftr* KO and WT mice (Figure 3C). Heatmap evaluation of the fifty most differentially abundant amplicon sequence variants (ASVs) reflected the differences of β-diversity between the iCftr KO and WT fecal samples (Figure 4). Six members of the genus *Muribaculaceae* were reduced in the iCftr KO versus WT feces. The family *Lachnospiraceae* had four members enriched in the iCftr KO samples and eight members significantly reduced relative to WT. At the species level, the iCftr KO demonstrated enrichment of four *Clostridia* (*C. innocuum*, *C. aldenense*-now *Enterocloster aldenense*, *C. bolteae*, *C.* UCG-014) and *Akkermansia muciniphilia* relative to WT. Table 1 shows the statistical analysis with Benjamini-Hochberg correction for the 50 most differentially abundant taxa. Overall, the loss of Cftr solely in the intestinal epithelium of iCftr mice consuming the PEG dietary regimen was sufficient to produce intestinal dysbiosis as compared to their WT sex-matched littermates.

**Figure 3.**
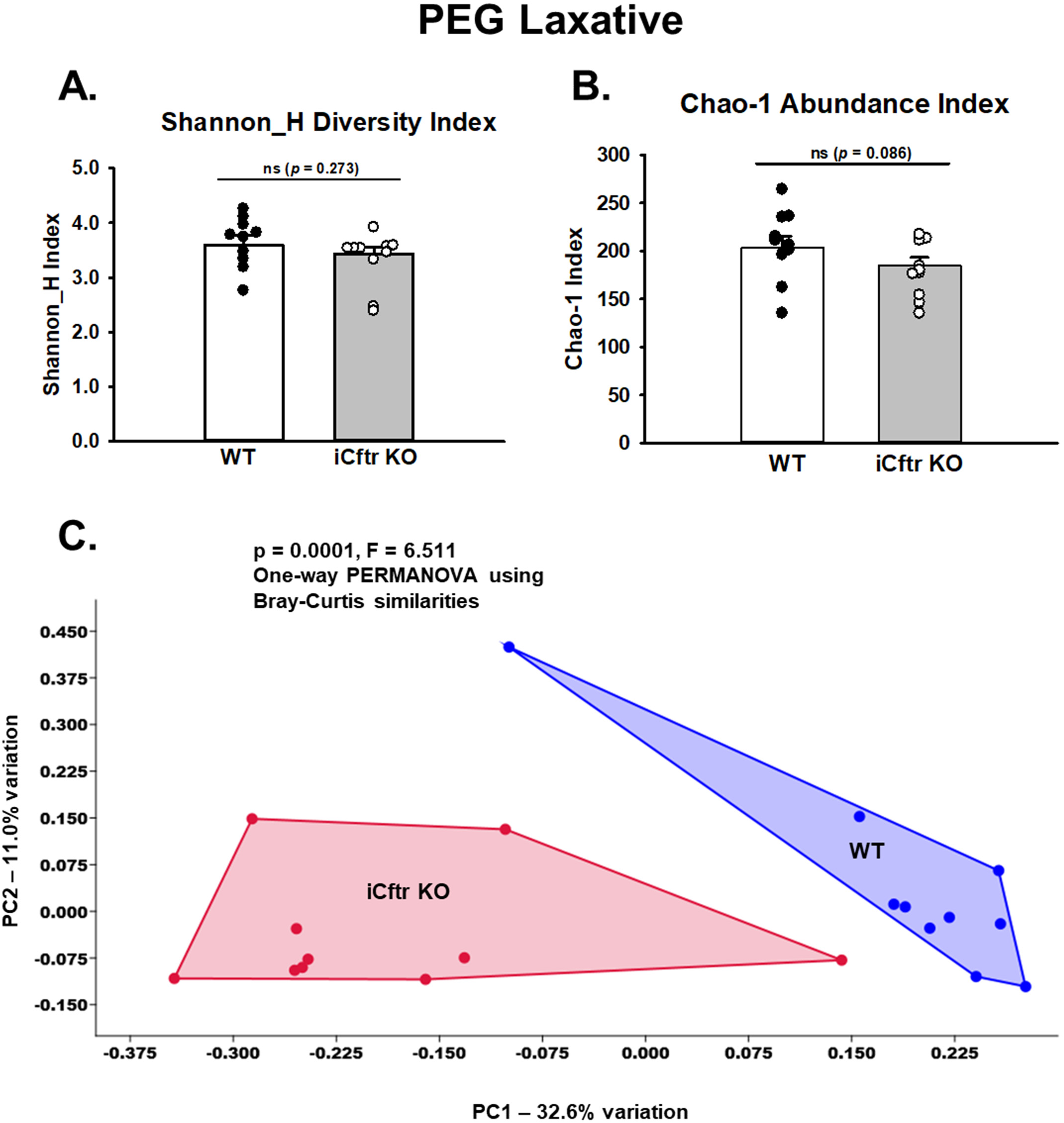
Alpha and beta diversities of the fecal microbiota from WT and iCftr KO sex-matched littermate pairs of mice receiving PEG laxative regimen. A.) Shannon H diversity Index of fecal microbial population evenness (p = 0.273). B.) Chao-1 abundance index of WT and iCftr KO mice receiving PEG laxative in drinking water (p = 0.086). Data represented as individual mouse values (WT – filled circles, iCftr KO – open circles) and mean ± SE (WT – white bars, iCftr KO – gray bars), ns = not significant via unpaired Student’s t-test. C.) Principal coordinate analysis plot based on the Bray-Curtis dissimilarity of the fecal microbial communities of WT (blue dots) and iCftr KO (red dots) mice receiving PEG laxative regimen. (p = 0.0001; F = 6.511). n = 10 WT and iCftr KO sex-matched littermate pairs, 6 male, 4 female.

**Figure 4.**
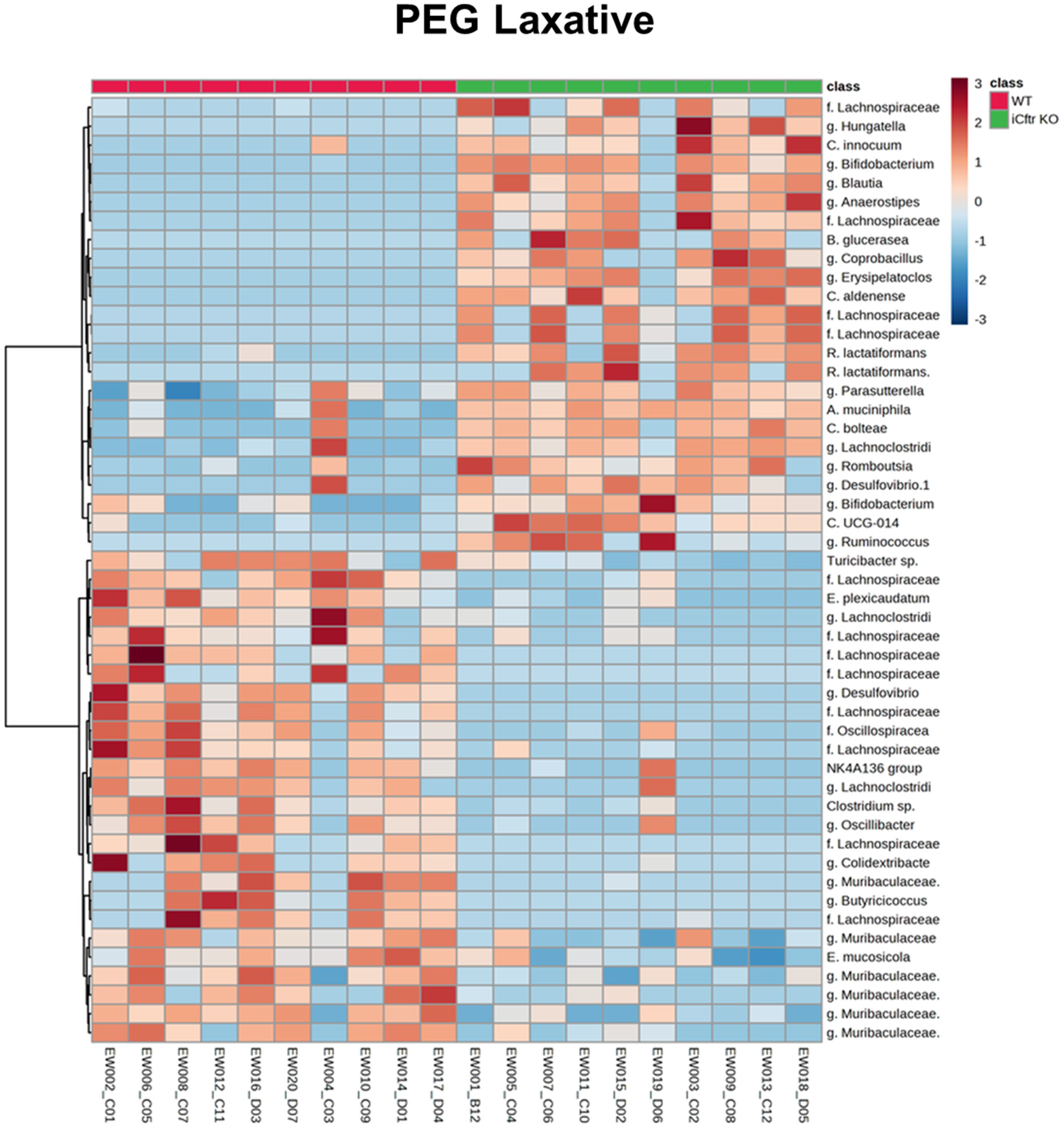
Heatmap of log transformed relative ASV abundance data of fecal samples from WT and iCftr KO sex-matched littermate mouse pairs receiving PEG laxative regimen. Samples (WT columns, red; iCftr KO columns, green) and 50 of the most differentially abundant OTUs (rows) arranged according to an unweighted pair group method with arithmetic mean (UPGMA) algorithm. n = 10 WT and iCftr KO sex-matched littermate pairs, 6 male, 4 female, legend upper right. f. = family, g. = genus.

**Table 1.**
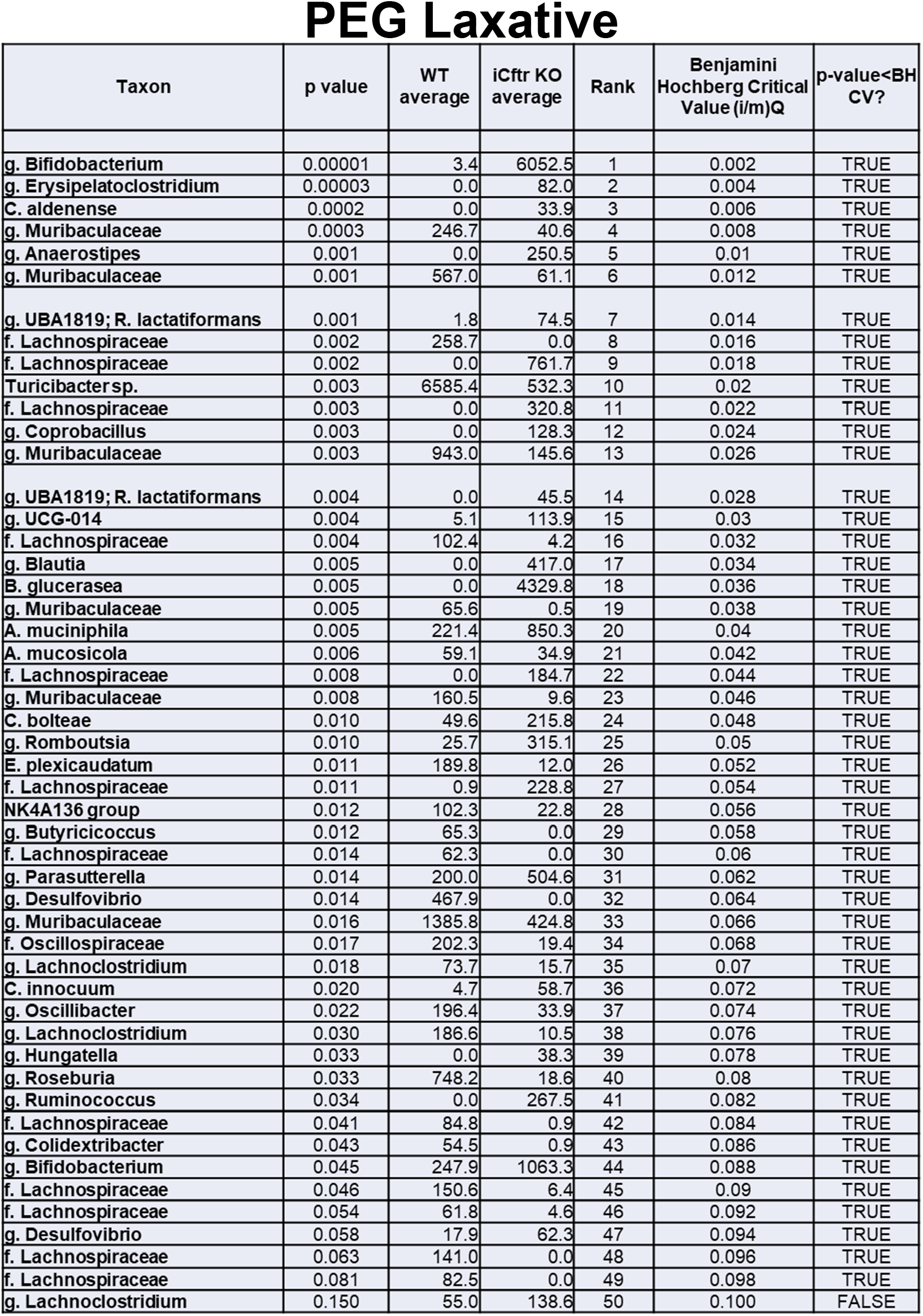
Table shows the 50 most differentially abundant fecal microbial taxa between WT and iCftr KO mice receiving PEG laxative regimen. p values were calculated using Student’s t-test, and significance determined using Benjamini Hochberg corrected p values. p < 0.05; m= 50, q = 0.1; n = 10 WT and iCftr KO sex-matched littermate pairs, 6 males, 4 females. f. = family, g. = genus.

### *iCftr* KO mice on the LiqD display significant dysbiosis with overt loss of bacterial species and decreases in Shannon diversity and Chao-1 abundance indices

To evaluate the effect of intestinal epithelial-specific loss of *Cftr* on the fecal microbiome in mice maintained on the LiqD dietary regimen, fecal samples were collected from i*Cftr* KO mice and WT sex-matched littermates after consuming the LiqD regimen for a 14-day period. Body weight and LiqD consumption was maintained or increased during the two-week period with all mice completing the regimen (data not shown). No significant effect of sex on the microbiome was detected, so data from five male and five female mouse pairs were combined for each group. Analysis of changes to relative abundance of bacterial phyla (Figure 5A) and bacterial families (Figure 5B) revealed a significant decrease in phyla including *Bacteroidota* (*p* < 0.01), *Desulfobacterota* (*p* < 0.01) and *Patescibacteria* (*p* < 0.05) in i*Cftr* KO fecal samples with an expansion of *Proteobacteria (Pseudomonadota)* (*p* < 0.01), as compared to WT. Looking at the family level in *Bacteroidota*, the i*Cftr* KO fecal samples showed significant increases in *Rikenellaceae* but more highly significant decreases in the families *Muribaculaceae* and *Tannerellaceae* (*p* < 0.01). Changes to families within the phyla *Patescibacteria* and the sulfate-reducing *Desulfobacterota* were not detected. The expansion of the phylum *Proteobacteria* (*Pseudomonadota*) included a significant increase in the family of *Enterobacteriaceae*. Many changes occurred in the *Firmicutes* phylum. Decreases in the families *Ruminococcaceae* (*p* < 0.05) and *Oscillospiraceae* (*p* < 0.001) were noted. On the other hand, other members of phylum *Firmicutes* were increased relative to WT, most notably the families *Clostridiaceae* and *Peptostreptococcaceae* (*p* < 0.05).

**Figure 5.**
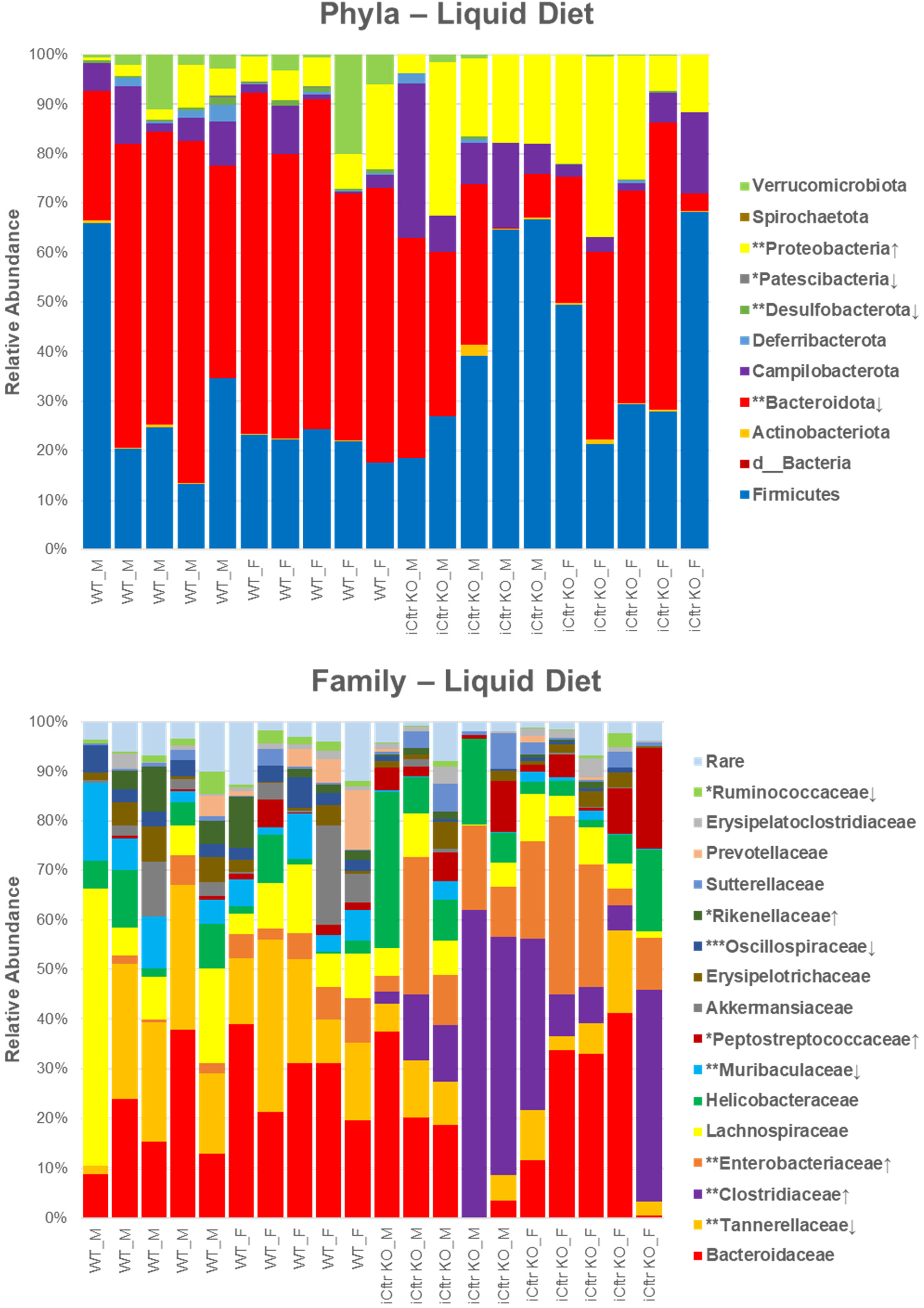
Bacterial taxa of fecal microbiome from WT and iCftr KO sex-matched littermate pair mice receiving LiqD regimen. Top. Stacked bar chart showing taxonomic relative abundance distribution of fecal microbial communities at the phylum level of WT and iCftr KO mice receiving the LiqD regimen. Bottom. Stacked bar chart showing taxonomic relative abundance distribution of fecal microbial communities at the family level of iCftr KO and WT mice on the LiqD regimen. Samples are grouped by genotype and sex at the x-axis. N = 10 WT and iCftr KO sex-matched littermate paired mice, 5 male, 5 female. Colored squares to the right of the charts indicate phylum or family identity. d = domain. ↑ = significantly increased relative to WT, ↓ = significantly decreased relative to WT. ***p-value<0.001, **p-value<0.01, *p-value<0.05 via unpaired Student’s t-test.

Regarding measures of α-diversity and richness, the Shannon diversity index (Figure 6A) and Chao-1 index (Figure 6B) both revealed significant decreases in iCftr KO fecal samples as compared to WT. As a measure of β-diversity, the PCoA revealed distinct clustering of the microbiota of iCftr KO and WT mice along the first principal coordinate (Figure 6C). Heatmap evaluation of the fifty most differentially abundant ASVs revealed overt differences of β-diversity in the iCftr KO versus WT fecal samples (Figure 7). Of the 11 taxa enriched in the iCftr KO versus WT samples, seven were members of the *Clostridium genus*, including *C. perfringens* (five strain/subtypes), *C. innocuum*, and the pathobiont *C. difficile*. The other four taxa with increased abundance in the iCftr KO samples were the genera *Erysipelatoclostridium*, *Tyzzerella* and *Helicobacter*, and the species *Ruminococcus gnavus.* Interestingly, in contrast to iCftr KO mice on the PEG regimen, the fecal samples from iCftr KO mice consuming the LiqD revealed a significant decrease of the mucolytic genus *Akkermansia*. Table 2 shows the statistical analysis with Benjamini-Hochberg correction for the 50 most differentially abundant taxa Overall, the loss of Cftr solely in the intestinal epithelium of iCftr mice consuming a LiqD dietary regimen produced a microbiome enriched with potential bacterial pathobionts.

**Figure 6.**
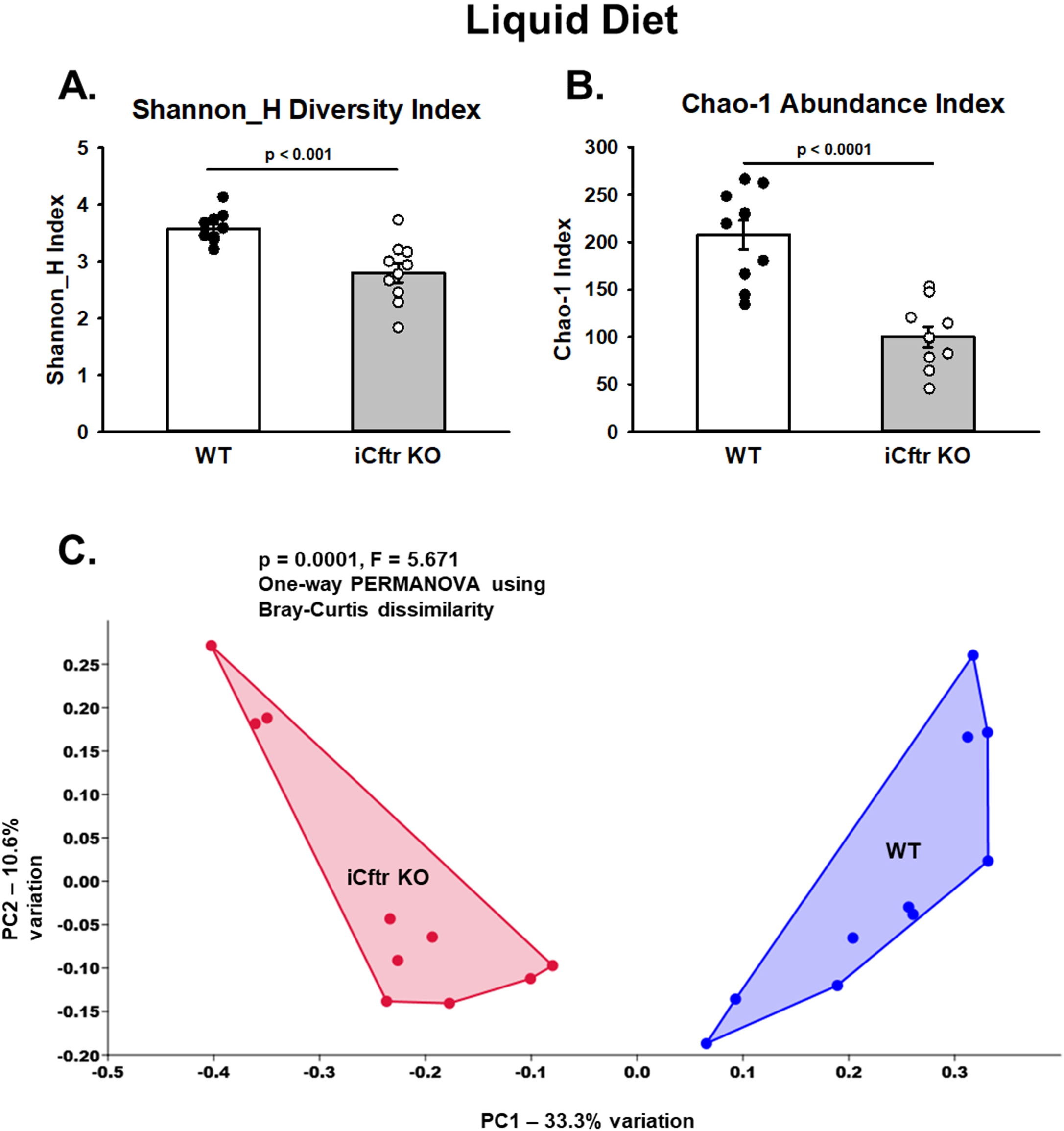
Alpha and beta diversities of the fecal microbiota from WT and iCftr KO sex-matched littermate pairs of mice receiving the LiqD regimen. A.) Shannon H diversity Index of fecal microbial population evenness (p < 0.001). B.) Chao-1 abundance index of WT and iCftr KO mice receiving the LiqD (p < 0.0001). Data represented as individual mouse values (WT – filled circles, iCftr KO – open circles) and mean ± SE (WT – white bars, iCftr KO – gray bars). C.) Principal coordinate analysis plot based on the Bray-Curtis dissimilarity of the fecal microbial communities of WT (blue dots) and iCftr KO (red dots) mice receiving the LiqD regimen. (p = 0.0001; F = 5.674). n = 10 WT and iCftr KO sex-matched littermate pairs, 5 male, 5 female.

**Figure 7.**
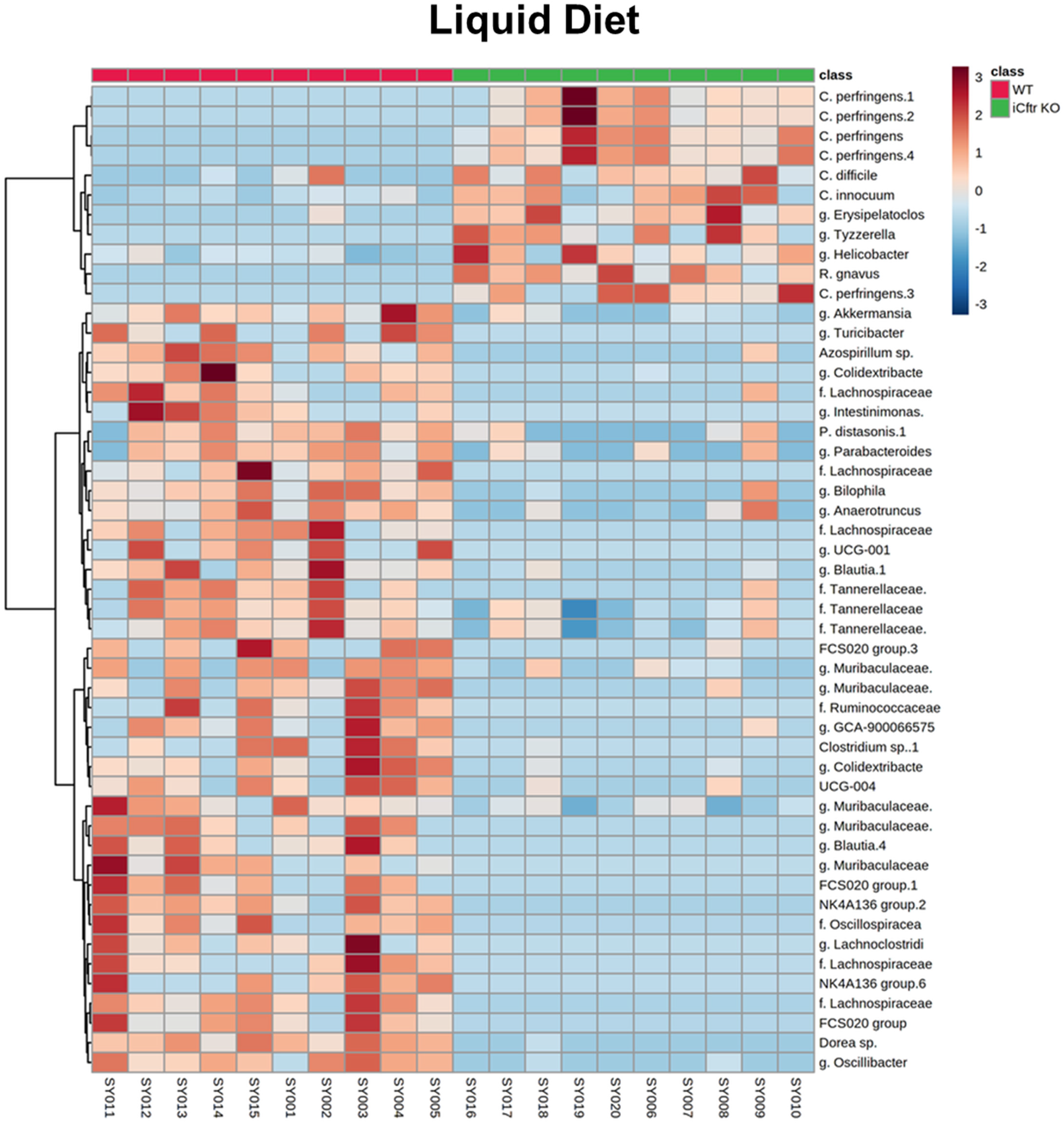
Heatmap of log transformed relative ASV abundance data of fecal samples from WT and iCftr KO sex-matched littermate mouse pairs receiving the LiqD regimen. Samples (WT columns, red; iCftr KO columns, green) and 50 of the most differentially abundant OTUs (rows) arranged according to an unweighted pair group method with arithmetic mean (UPGMA) algorithm. n = 10 WT and iCftr KO sex-matched littermate pairs, 5 male, 5 female, legend upper right. f. = family, g. = genus.

**Table 2.** Table shows the 50 most differentially abundant fecal microbial taxa between WT and iCftr KO mice receiving the LiqD regimen. p values were calculated using Student’s t-test, and significance determined using Benjamini Hochberg corrected p values. p < 0.05; m= 50, q = 0.1; n = 10 WT and iCftr KO sex-matched littermate pairs, 5 males, 5 females. f. = family, g. = genus.

Liquid Diet
| Taxon | t-test (WT vs. iCftr KO) | WT Average | iCftr KO Average | Rank | Benjamini Hochberg Critical Value | p-value < BH CV? |
| --- | --- | --- | --- | --- | --- | --- |
| Dorea sp. | 0.0001 | 100.6 | 0.7 | 1 | 0.002 | TRUE |
| g. Oscillibacter | 0.0003 | 447.5 | 5.1 | 2 | 0.004 | TRUE |
| NK4A136 group | 0.0008 | 81.5 | 0.0 | 3 | 0.006 | TRUE |
| f. Lachnospiraceae | 0.001 | 48.0 | 0.0 | 4 | 0.008 | TRUE |
| g. Muribaculaceae | 0.001 | 217.9 | 24.6 | 5 | 0.010 | TRUE |
| g. Muribaculaceae | 0.002 | 77.6 | 0.0 | 6 | 0.012 | TRUE |
| g. Parabacteroides | 0.003 | 144.9 | 40.2 | 7 | 0.014 | TRUE |
| P. distasonis | 0.003 | 1543.5 | 427.1 | 8 | 0.016 | TRUE |
| g. Muribaculaceae | 0.004 | 200.1 | 11.6 | 9 | 0.018 | TRUE |
| R. gnavus | 0.004 | 0.0 | 754.7 | 10 | 0.020 | TRUE |
| g. UCG-004 | 0.005 | 134.5 | 13.4 | 11 | 0.022 | TRUE |
| Azospirillum sp. | 0.007 | 179.9 | 9.1 | 12 | 0.024 | TRUE |
| C. perfringens | 0.007 | 0.0 | 15307.4 | 13 | 0.026 | TRUE |
| C. innocuum | 0.007 | 26.3 | 856.4 | 14 | 0.028 | TRUE |
| g. Turicibacter | 0.008 | 36.4 | 0.0 | 15 | 0.030 | TRUE |
| f. Tannerellaceae | 0.008 | 109.1 | 9.4 | 16 | 0.032 | TRUE |
| C. perfringens | 0.008 | 0.0 | 429.0 | 17 | 0.034 | TRUE |
| NK4A136 group | 0.009 | 26.4 | 0.0 | 18 | 0.036 | TRUE |
| f. Oscillospiraceae | 0.009 | 145.7 | 0.0 | 19 | 0.038 | TRUE |
| C. perfringens | 0.010 | 0.0 | 808.0 | 20 | 0.040 | TRUE |
| g. Tyzzerella | 0.011 | 0.0 | 638.8 | 21 | 0.042 | TRUE |
| g. Bilophila | 0.011 | 254.4 | 38.2 | 22 | 0.044 | TRUE |
| f. Tannerellaceae | 0.012 | 3883.2 | 1159.0 | 23 | 0.046 | TRUE |
| FCS020 group | 0.012 | 195.8 | 0.0 | 24 | 0.048 | TRUE |
| f. Lachnospiraceae | 0.012 | 183.5 | 0.0 | 25 | 0.050 | TRUE |
| g. Lachnospiraceae FCS020 | 0.013 | 55.9 | 0.0 | 26 | 0.052 | TRUE |
| g. UCG-001 | 0.019 | 216.1 | 0.0 | 27 | 0.054 | TRUE |
| g. Muribaculaceae | 0.020 | 230.4 | 32.1 | 28 | 0.056 | TRUE |
| f. Ruminococcaceae | 0.021 | 60.5 | 0.0 | 29 | 0.058 | TRUE |
| g. Blautia | 0.022 | 59.3 | 0.0 | 30 | 0.060 | TRUE |
| g. GCA-900066575 | 0.022 | 56.8 | 2.0 | 31 | 0.062 | TRUE |
| g. Lachnospiraceae FCS020 group | 0.024 | 17.3 | 0.5 | 32 | 0.064 | TRUE |
| Clostridium sp. | 0.025 | 37.9 | 0.3 | 33 | 0.066 | TRUE |
| f. Lachnospiraceae | 0.027 | 31.6 | 2.9 | 34 | 0.068 | TRUE |
| f. Lachnospiraceae | 0.031 | 15.4 | 0.0 | 35 | 0.070 | TRUE |
| g. Erysipelatoclostridium | 0.032 | 11.1 | 534.5 | 36 | 0.072 | TRUE |
| g. Colidextribacter | 0.032 | 143.0 | 3.2 | 37 | 0.074 | TRUE |
| g. Intestinimonas | 0.033 | 38.6 | 0.0 | 38 | 0.076 | TRUE |
| g. Helicobacter | 0.036 | 294.1 | 4170.1 | 39 | 0.078 | TRUE |
| f. Tannerellaceae | 0.038 | 656.6 | 201.6 | 40 | 0.080 | TRUE |
| g. Akkermansia | 0.052 | 3898.5 | 237.2 | 41 | 0.082 | TRUE |
| g. Muribaculaceae | 0.061 | 783.8 | 0.0 | 42 | 0.084 | TRUE |
| g. Lachnoclostridium | 0.068 | 106.0 | 0.0 | 43 | 0.086 | TRUE |
| g. Blautia | 0.070 | 523.9 | 10.5 | 44 | 0.088 | TRUE |
| g. Anaerotruncus | 0.075 | 773.8 | 215.2 | 45 | 0.090 | TRUE |
| C. difficile | 0.083 | 455.7 | 1833.4 | 46 | 0.092 | TRUE |
| C. perfringens | 0.084 | 0.0 | 1166.9 | 47 | 0.094 | TRUE |
| f. Lachnospiraceae | 0.088 | 260.4 | 0.0 | 48 | 0.096 | TRUE |
| C. perfringens | 0.096 | 0.0 | 911.3 | 49 | 0.098 | TRUE |
| g. Colidextribacter | 0.154 | 187.4 | 0.4 | 50 | 0.100 | FALSE |

## Discussion

Increasing research on the fecal microbiome of children with CF indicates dysbiosis of immune-modulating bacteria can affect local and systemic immunity early in life and, thereby, the development of a potentially unfavorable gut-lung axis(8, 40–43). The intestinal manifestations of CF extend well into adulthood with chronic bowel inflammation and an increased risk of gastrointestinal cancer. Similar to inflammatory bowel disease(44), significant intestinal dysbiosis is associated with bowel inflammation in CF patients(41). CFTR is expressed in multiple organs and tissues, which may influence the intestinal microbiota, as has been shown by studies of myeloid cell Cftr KO mice(29, 30, 32, 45, 46). To extend studies of tissue-dependent effects on dysbiosis in the CF intestine, the present study evaluated epithelial-specific deletion of Cftr in the intestine. The iCftr KO mice exhibited intestinal inflammation and overt dysbiosis that had significant similarities to previous studies of pan-Cftr KO mice. The pan-Cftr KO mouse accentuates the pathology of the human CF intestinal disease with high morbidity from meconium ileus-like disease that requires the constant provision of either PEG or LiqD, throughout life to enable the utility of the animal model(24–26). Therefore, both dietary interventions were investigated to inform on the gut environment for on-going or future studies employing CF mouse models.

### PEG laxative

Intestinal-specific Cftr KO mice consuming PEG laxative regimen displayed less overt features of inflammation and fecal dysbiosis, as compared to the iCftr KO mice on the LiqD. Fecal calprotectin measurement indicated greater inflammation was present in the iCftr KO relative to WT. Transcriptomic studies of pan-Cftr KO mice maintained on the PEG laxative also show slight changes in inflammatory markers with increases in only resistin-like molecule B and hematopoietic cell transcript(38). The mild nature of dysbiosis was indicated by no change in α-diversity of the PEG iCftr KO microbiome relative to WT, which aligns favorably with one study of the small bowel microbiome of C57BL/6J pan-Cftr KO mice(15). However, other studies of PEG pan-Cftr KO microbiota of small bowel and large bowel have found changes in α-diversity(14, 17, 32). Like the iCftr KO analysis, most studies of pan-Cftr KO microbiota have found a highly significant difference in β-diversity (compositional similarity) clustering to WT(14, 15, 17, 32).

Alterations in the bacterial phyla of the fecal microbiome in PEG-treated iCftr KO mice relative to WT were similar to increases in *Actinobacteriota* (*Actinomycetota*), *Verrucomicrobiota* and *Proteobacter* (*Pseudomonadota*) of the small intestinal microbiome in pan-Cftr KO mice (C57BL/6J strain)(15). Within the *Actinobacteria* phylum, the iCftr KO fecal microbiome and studies of PEG pan-Cftr KO samples consistently show significant increases in genera of *Bifidobacteriaceae*, which are considered beneficial symbionts(14, 15, 17, 32, 47, 48). Increases in *Verrucomicrobiota* phylum in the PEG iCftr KO involved increases in *Akkermansiaceae*, a known mucus-degrading bacteria whose population is inversely related to intestinal inflammation(49), which is consistent with one study of pan-Cftr KO small intestinal microbiome but not others (14, 15, 17, 32). Increases in *Proteobacteria* (*Pseudomonadota*) phylum in the PEG iCftr KO fecal microbiome, known to be associated with IBD and metabolic disease(50–52), is consistent with other studies of pan-Cftr KO microbiota but with differences in taxa(14, 17). Similarly, decreases in the phyla *Bacteriodota* and *Desulfobacterota* in the PEG iCftr KO fecal samples were found in studies of pan-Cftr KO microbiota but with differences in taxa (17, 32).

Although *Firmicutes* (*Bacillota*) was unchanged in the PEG iCftr KO samples, there were significant changes (increases and decreases) in several taxa. One study of pan-Cftr KO mice found a decrease in the phylum *Firmicutes*(15). Notably, both studies had significant decreases in *Erysipelotrichaceae*, a change associated with Crohn’s disease(53). Other studies of pan-Cftr KO microbiomes had similar offsetting changes to taxa within *Firmicutes*(14, 32). Analysis of the iCftr KO most abundant taxa indicated enrichment of four *Clostridium* species, including *C. innocuum*, which has characteristics consistent with *C. difficile*(54) and increases in *Peptostreptococcaceae*, which has been associated with IBD and colorectal cancer(55, 56). Overall, the microbial changes in the iCftr KO fecal microbiome of mice maintained on the PEG laxative show a mild inflammatory state and dysbiosis with features similar to pan-Cftr KO intestine.

### Liquid Diet

An advanced disease phenotype was predicted from the provision of the LiqD regimen based on previous studies by De Lisle et al.(16, 28, 38). This was confirmed by a significant increase in fecal calprotectin relative to WT in the iCftr KO mice consuming the LiqD. Significant decreases in α- and β-diversity were exhibited by the iCftr KO fecal microbiota mice, which matches with a study of the LiqD pan-Cftr KO ileum(16). A major finding of the iCftr KO microbiota was the enrichment of phylum *Proteobacteria* (*Pseudomonadota*) family *Enterobacteriaceae* and phylum *Firmicutes* families *Clostridiaceae* and *Peptostreptococcaceae* when consuming the LiqD. These families are known to include several pathobionts (55–57). Further, previous studies of LiqD pan-Cftr KO ileal microbiota found evidence that these families may suppress bacterial diversity and gut-homeostatic bacteria in comparisons before and after a 3-week antibiotic regimen (ciprofloxacin, metronidazole)(16). In line with this observation, the LiqD iCftr KO fecal samples showed decreases in the phyla *Bacteriodota*, *Patescibacteria*, *Desulfobacterota* and beneficial *Firmicutes* families, *Ruminococcaceae* and *Oscillospiraceae*, both are butyrate producers and decreases in *Ruminococcaceae* has been associated with IBD(58, 59). It was surprising that among the 11 most differentially abundant species in the iCftr KO fecal samples, seven were *Clostridial* species with *C. perfringens* (5 subtypes/subvariants), *C. difficile* and *C. innocuum*, which are associated with toxin production and disease(54, 60, 61). *C. difficile* is seen in the stool of approximately 50% of sampled CF patients and is found to mainly produce Toxin B(62). Taxa with increased abundance in the iCftr KO samples included *Erysipelatoclostridium*, *Tyzzerella* and *Helicobacter*, which also have known associations with IBD(63–65). Assessment of the 50 most differentially abundant taxa revealed mucolytic *Ruminococcus gnavus* in the LiqD iCftr KO feces, which is associated with Crohn’s disease in strains that elaborate an inflammatory polysaccharide toxin(66). In contrast, decreases in the beneficial genus *Akkermansia* were found in the LiqD iCftr KO relative to WT. Differences in mucolytic species in iCftr KO fecal samples on mice consuming LiqD versus PEG is likely dependent on multiple factors. Among these, it is possible that intestinal mucus is more tenacious and concentrated in iCftr KO mice on the LiqD as compared to those on the PEG laxative which is high in osmolytes and contains HCO_3_ (∼11 mM/L) that increases the solubility of Cftr KO mucus(67). Although intestinal dysbiosis and inflammation were more severe in iCftr KO mice consuming the LiqD, both dietary regimens share a common feature. Studies performed on pan-Cftr KO small intestine on either dietary regimen have been shown to exhibit mucus casts within the crypts that entrap secreted Paneth cell granules, thereby, limiting elaboration of anti-microbials (lysozyme, α-defensins) into the intestinal lumen(12, 68). This is a driver of dysbiosis akin to Paneth cell dysfunction in Crohn’s disease(69). Translationally, PEG laxative use is not routinely prescribed for pwCF, but the present study suggests that low level daily use might yield benefits with regard to intestinal dysbiosis.

The limitations of the present study are several. As mentioned, study of the intestinal-specific Cftr KO fecal microbiome was designed to determine the effect of two anti-obstructive dietary regimens, but these regimens differ in dietary composition, in particular, fiber content. Inclusion of additional fermentable fiber with the LiqD may have prevented some of the dysbiosis and inflammation through the generation of intraluminal short-chain fatty acids(41, 70). The two-week switch to the LiqD was based on the time required for microbiota to stabilize after transfer(71), but a longer time period may be necessary to fully ‘mature’ the microbiota. The WT and iCftr KO sex-matched littermates were co-housed until weaning but were individually housed during the experimental period. This study design does not provide the influence of co-mingling genotypes, but it did demonstrate that the development of the microbiome in individual WT and iCftr mice is surprisingly consistent.

Dysbiosis and associated inflammation are recognized to have untoward effects on the general health and well-being of pwCF(42). Our studies indicate that intestinal dysbiosis and bowel inflammation is largely driven by the loss of Cftr in the intestinal epithelium, but varies in magnitude dependent on the anti-obstructive measures employed. The Cftr KO mice provide the opportunity to investigate CF intestinal disorders in a controlled model that is not confounded by co-morbidities or differing diets and therapeutic measures that occur in studies of pwCF. However, studies of Cftr KO intestine or mucosal/systemic immunity will need to consider the impact of anti-obstructive interventions on the gut microbiota in the experimental design.

## DATA AVAILABILITY

All sequence data supporting the current manuscript are openly available in the National Center for Biotechnology Information (NCBI) Sequence Read Archive (SRA, https://www.ncbi.nlm.nih.gov/sra) as BioProject ID: PRJNA993973.

## GRANTS

Funding for this research was provided by a University of Missouri College of Veterinary Medicine Phi Zeta research grant (Young/Clarke), National Institutes of Health CTSA Grant Number UL1TR002345 (Clarke), Cystic Fibrosis Foundation grant CLARKE20G0 (Clarke) and Crohn’s and Colitis Foundation 693938 (Clarke).

## AUTHOR CONTRIBUTIONS

S.M.Y., R.A.W., and L.L.C. conceived and designed research; S.M.Y, R.A.W., E.W., and A.E. performed experiments; S.M.Y., R.A.W., E.W., A.E., and L.L.C. analyzed data; S.M.Y., R.A.W., E.W., A.E., and L.L.C. interpreted results of experiments; S.M.Y., R.A.W., E.W., and L.L.C. prepared figures; S.M.Y., R.A.W. and L.L.C. drafted manuscript; S.M.Y., R.A.W., A.E., and L.L.C edited and revised manuscript; S.M.Y., R.A.W., E.W., A.E., and L.L.C approved final version of manuscript.

## ACKNOWLEDGMENTS

The authors wish to acknowledge the University of Missouri Metagenomics Center, the University of Missouri Genomics Technology Core and the University of Missouri Bioinformatics and Analytics Research Core for their assistance in sample processing and data formation. We thank Dr. Craig Franklin and Dr. James Amos-Landgraf (University of Missouri) for their assistance with reviewing the prepared manuscript. We would also like to thank Dr. Craig Hodges (Case Western Reserve University) for assisting us with establishing the intestinal-specific *Cftr* knockout colony at our institution.

The present address of S. M. Young: 1425 Plain-City Georgesville Rd, Building JM 10, West Jefferson, OH 43162.

